# Functional fingerprinting for the developing brain using deep metric learning: an ABCD study

**DOI:** 10.1101/2024.12.13.628288

**Authors:** Rui Xu, Shuwan Zhao, Qianhui Jin, Zeyao Wang, Kun Zhao, Suyu Zhong, Jiaying Zhang, Yong Liu, Ting Qi, Yongbin Wei

## Abstract

Brain fingerprinting is a promising approach for characterizing the uniqueness of individual brain functioning using functional magnetic resonance imaging (fMRI) data. Here, we propose a novel deep learning framework, the metric-BoIT, for brain fingerprinting and demonstrate its effectiveness in capturing individual variability among early adolescents undergoing dramatic brain changes. Utilizing resting-state fMRI data from the Adolescent Brain Cognitive Development (ABCD) dataset, we identified brain functional fingerprints that achieved remarkable individual identification accuracy, reaching 97.6% within a single session and maintaining 86.6% accuracy over a four-year developmental period. Annotation analysis revealed that higher-order association regions, particularly those within the default-mode network, contributed most significantly to these distinctive brain fingerprints (*t* = 5.618, *p* < 0.001). Moreover, these brain fingerprints were relevant to cognitive functions, as evidenced by significant correlations with fluid intelligence (*F* = 1.282, *p* = 0.027) and crystallized intelligence (*F* = 1.405, *p* < 0.001). The extracted brain fingerprints were additionally associated with genetics, showing that individuals with strong genomic relationships exhibited more similar brain fingerprint patterns (*t* = −12.330, *p* < 0.001). Together, our study not only presents an innovative approach to brain functional fingerprinting but also provides valuable insights into the individual variability underlying adolescent neurodevelopment.

## INTRODUCTION

Individual differences in both the function and structure of the brain have been widely observed in adults (*1–5*) and are particularly pronounced during development (*6*, *7*). A considerable variation in the functional connectivity (FC) of association regions, in contrast to a relatively limited variation in primary sensorimotor and visual regions, has been noted as early as the third trimester of brain development (*8*, *9*). Similar inter-individual variation patterns of FC have also been reported in later development stages of childhood, with several higher-order brain networks further correlating with variations in children’s cognitive abilities (*10*). Characterizing the unique pattern of human brain development is thus critical for understanding the complex developmental trajectories underlying children’s behavioral and neurocognitive changes.

In light of the individual variability, researchers have introduced the “brain fingerprint”, which refers to distinctive neural signatures that enable the identification of individuals within a population (*11*). Brain fingerprint was proposed as a promising approach to characterize the uniqueness of brain functioning based on functional magnetic resonance imaging (fMRI) data. Using the functional brain connectome, Finn et al. precisely identified specific individuals from a large cohort of young adults and distinguished different brain activation patterns across tasks (*11*). These brain fingerprints rest upon FC matrices and exhibit stable and significant individual differences in FC distributed across three higher-order cognitive networks (i.e., default-mode, dorsal attention, and fronto-parietal networks) reflecting inherent functional dynamics of an individual’s brain connectome (*12*). The accuracy of individual identification for young adults, either using brain fingerprints of statical or dynamical FC, remains stable over several months (*13*), suggesting that the intrinsic individual connectivity patterns do not significantly change in adults. Moreover, brain fingerprints based on the functional connectome have demonstrated the capability of predicting cognitive traits or clinical symptoms in mental conditions (*14*, *15*). These findings stress that fingerprinting based on the functional connectome offers a straightforward and reliable approach for the examination of individual variability.

While brain fingerprints robustly identify the unique individual among adults, it is a different scenario in the context of developing populations given the dynamic nature of the brain during maturation. Using the functional connectome, a relatively lower accuracy has been observed when identifying individuals among children and adolescents compared to that of adults, indicating that individuals’ functional connectome continues to become unique during development (*16*). Moreover, longitudinal studies have shown that brain fingerprinting remained valid over periods of 3 months to 2 years in adolescent populations, however the accuracy of identification significantly dropped to below 80% compared to that of adults (usually over 90%) (*17*, *18*). Novel fingerprinting methods are imperative for identifying more refined developmental patterns of brain functioning, specifically functional fingerprints that could effectively differentiate individuals during a longer period of development.

Currently, various methods have been developed for extracting brain fingerprints using fMRI data. Finn et al. utilized a FC-based correlation approach, with individuals identified by calculating the correlation between two functional connectomes (*11*). Recent advances have incorporated brain dynamics through time-windowed analyses, segmenting time series into different windows to identify when the best identification occurs and at what time scale (*19*). Although these methods have made significant progress in identifying individuals, there are still some challenges. Notably, the identification accuracy using FC decreases significantly as the number of subjects increases, with an estimated accuracy of 62% for a sample size of 10,000 and 42% for 100,000 subjects (*20*). More recently, deep learning methods have further opened up new avenues for brain fingerprinting by maximizing inter-subject variances in embedding space (*21–25*). For instance, an effective approach combines autoencoders or conditional variational autoencoders with sparse dictionary learning modules to capture shared information among subjects, while using the residuals with enhanced inter-individual variability as brain fingerprints for individual identification (*26*, *27*). These methods demonstrate great potential in capturing the unique functional fingerprints of individual brain networks, yet more refined methodologies with enhanced identification precision during development as well as more solid neurobiological explainability are needed.

In the current study, we propose a deep learning framework based on distance metric learning for longitudinal identification of brain fingerprints in early adolescence. Utilizing the blood-oxygen-level-dependent transformer (BolT) (*28*), this framework optimizes the feature space by maximizing the inter-subject distances while minimizing intra-subject distances. Using longitudinal resting-state fMRI data from the ABCD dataset of children aged 9-10 years at baseline, we achieve high identification accuracies of 97.6% within one scan session and of 86.6% over four years period. Further, we demonstrate key regions within higher-order cognitive networks with significant individual variability during development and highlight the cognitive correlates and genetic influences on the brain fingerprints.

## RESULTS

### The framework of Metric-BolT

To extract brain fingerprints during brain development, we integrate the BoIT model (*28*) with deep metric learning, further referred to as “Metric-BolT” (see MATERIALS AND METHODS for details). We applied Metric-BoIT to resting-state fMRI data from the ABCD dataset, including fMRI time series data from a total of 1325 individuals scanned at three time points (i.e., baseline, two and four years later). Applying Metric-BolT on the longitudinal dataset allows us to perform individual identification across different time points and intervals, providing insights into investigating individual variability in functional dynamics throughout adolescence.

The detailed framework of Metric-BolT is shown in Fig. 1. During training the model processes three data matrices of time series, with two from the same subject at different runs (e.g., baseline and two years later), and one from a different subject (e.g., either baseline or two years later). The BoIT transformer blocks yield distinct feature vectors, referred to as brain fingerprints, for each data matrix. A tripletMarginLoss function is used to minimize the distance between feature vectors of the same subject while maximizing the distance between vectors from different subjects. Once the model was trained, a database of brain fingerprints was generated by applying the trained Metric-BoIT to the one run of testing samples. Given a target subject from another run of testing samples, individual identification was accomplished by matching the target sample to the fingerprint from the database with the closest distance (Fig. 2A).

**Fig. 1.**
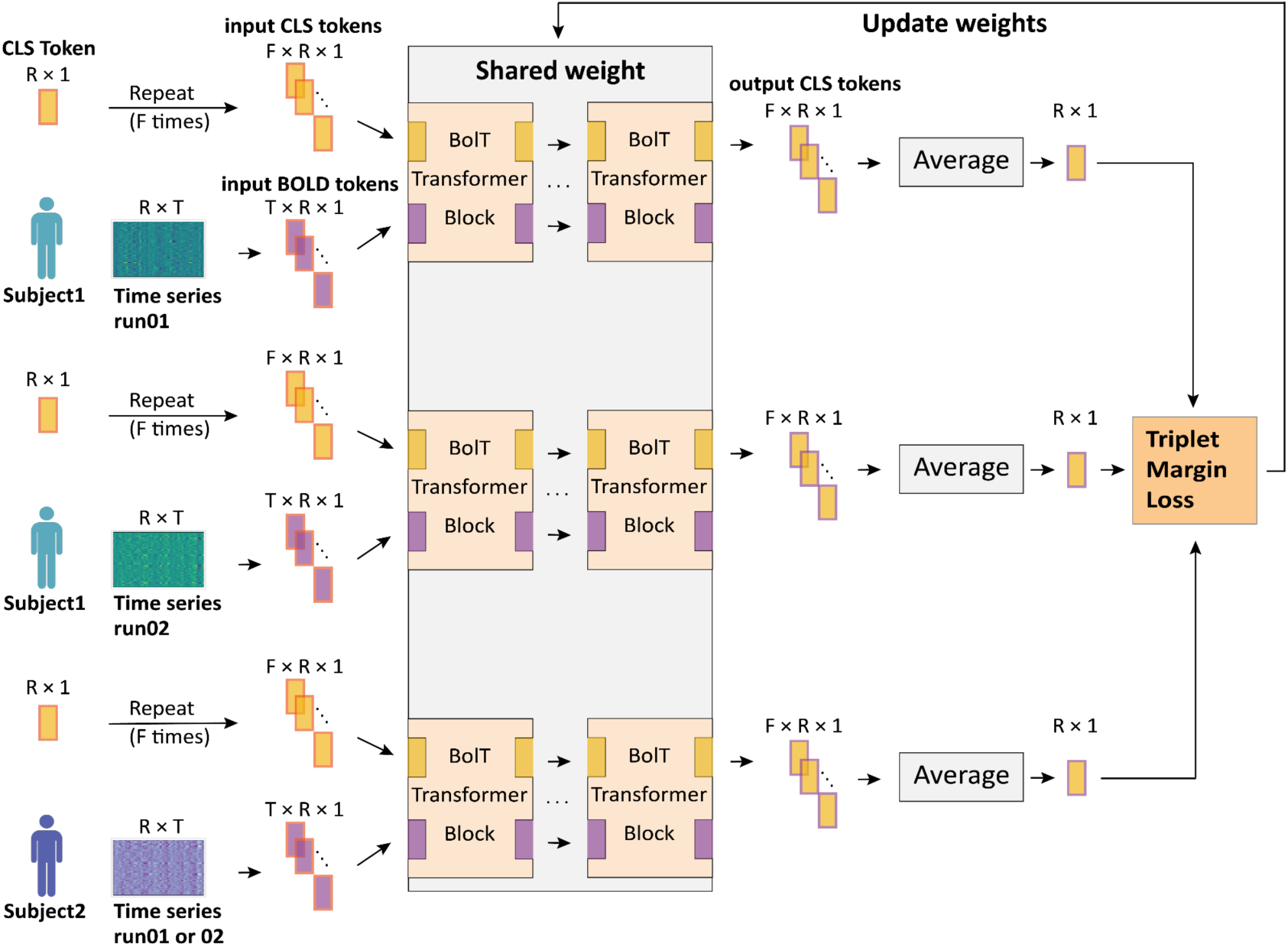
The scheme of Metric-BolT. Each training iteration involves three time series as inputs: two from the same subject (as the anchor and positive sample) and one from a different subject (the negative sample). Each time series matrix is divided into BOLD tokens, then processed through transformer blocks. Within each transformer block, the time series is further divided into multiple temporally overlapping windows, with each window containing multiple BOLD tokens. CLS tokens (see MATERIALS AND METHODS) are initialized for each time window to capture window-specific information. The averaged output CLS tokens represent the global features of the original input, forming the feature vectors (i.e., brain fingerprints). The model is trained and updated using TripletMarginLoss as its optimization criterion. Notably, all three samples in a single training iteration share the same model weights. R: number of brain regions; T: time series length; F: number of time windows.

**Fig. 2.**
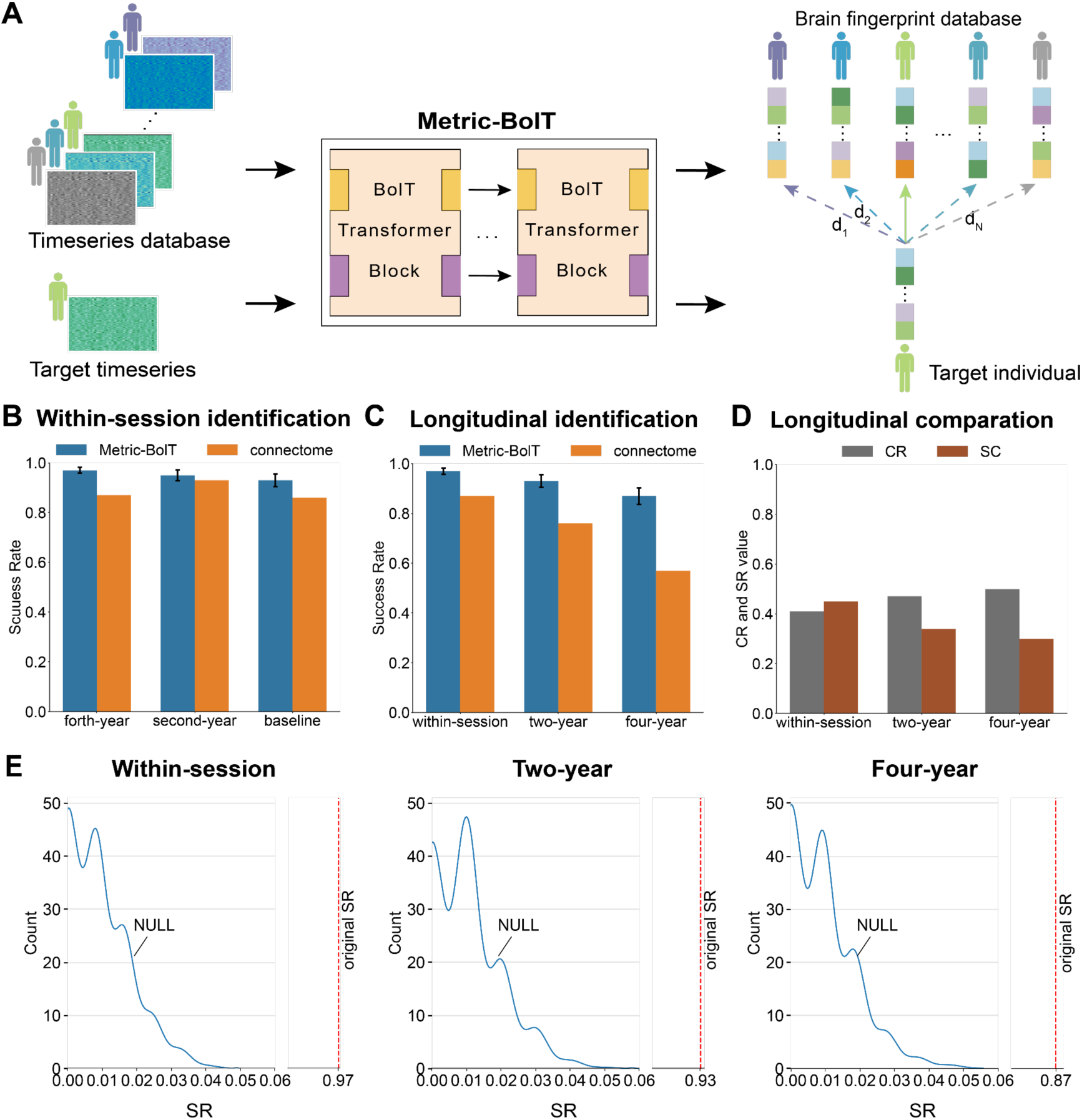
Individual identification results across various time intervals. (A) Given a target data sample, we transformed the database time series and the target time series into database feature vectors and target feature vectors using the Metric-BolT model. We then computed the distances between the target vector and each vector in the database, identifying the vector with the minimum distance. (B) SR of individual identification using Metric-BolT and connectome-based approach at baseline, second year follow-up, and fourth year follow-up data. (C) SR of individual identification using metric-BolT and connectome-based approach at within-session (two runs from fourth year follow-up), two-year (runs from fourth and second year follow-up), and four-year data intervals. (D) The ratio of average intra-class to inter-class distances (CR) and the silhouette coefficients (SC) for different time intervals. (E) The distribution of SR values in permutation tests, where the x axis represents SR values and the y axis, Count, represents the frequency of each SR value. The solid blue line indicates the actual distribution of SR values, while the red dashed line represents the SR under the non-permutation condition.

### Metric-BolT precisely identifies individuals longitudinally

We started by evaluating the performance of Metric-BolT in identifying individuals across three time points during brain development. To assess the overall accuracy of the identification, success rate (SR) that measures the proportion of correctly identified individuals was computed (see MATERIALS AND METHODS for details). Individual identification was first conducted using data from the two runs collected within the same scan session, taking data samples from the first run as the fingerprint database and testing each data sample from the second run (as target samples) individually. Metric-BolT achieved a SR of 93.4% for the two runs of baseline data, 95.2% for the second-year follow-up data, and 97.3% for the fourth-year follow-up data (Fig. 2B). This result confirmed an increasing distinctiveness of the functional fingerprints along with brain development. Moreover, longitudinal identification, taking data from a baseline run as the fingerprint database and testing individuals from a second-year run, showed an SR of 90.9%, while testing the four-year run showed an SR of 86.6%. Testing the fourth-year individuals against the second-year fingerprint database further revealed an SR of 92.6%. Reversing the roles of the brain fingerprint database and testing samples revealed similar results: within-session runs showed an SR of 94.0%, 96.3%, 97.6% at the baseline, second-year follow-up, and fourth-year follow-up, respectively. Swapping the baseline run and second-year run, the second-year run and fourth-year run, and the baseline run and fourth-year run respectively showed an SR of 91.6%, 93.0%, and 86.6% (Fig. 2C). These high SRs in individual identification indicate that the identified brain fingerprints are effective and maintain relatively stable over an interval of even four years during development. Additionally, the achieved SR outperforms the SR obtained using the functional connectome-based approach (*11*) (Fig. 2B and Fig. 2C).

To assess the statistical significance of identification accuracy, we conducted permutation tests to examine whether the achieved SRs were greater than by chance (see details in MATERIALS AND METHODS). The maximum SRs were 4.07% for identifications within one session, 5.00% for the two-year interval, and 4.59% for the four-year interval, obtained under random permutations (all *p* < 0.001, 1000 permutations) (Fig. 2E). These results confirmed the achieved SRs to be higher than what can be expected by chance.

In addition to SR, we used averaged intra- and inter-class distance ratio (CR) and the Silhouette coefficient (SC) to evaluate the individual identification performance. CR quantifies the separation of feature vectors by comparing distances within the same subject to those between different subjects. SC evaluates how similar each feature vector is to other vectors from the same subject compared to those from the nearest different subject (see MATERIALS AND METHODS for details). CR rose from 0.41 for within-session individual identification to 0.47 for identification with a two-year interval, and to 0.5 for identification with a four-year interval (Fig. 2D). Meanwhile, SC dropped from 0.45 to 0.34 and then to 0.30 along with the time intervals (Fig. 2D), also suggesting that a higher individual identification performance is associated with a shorter time interval.

### Brain fingerprint is driven by brain association regions

To understand the key factors driving brain fingerprints, we next examined the importance of each brain region during brain fingerprint extraction, using regional weights (*W*) from random forest models to represent the relative contribution of each brain region, with a larger weight *W* indicating higher contributions (see also MATERIALS AND METHODS). This quantified the individual variations in functional dynamics of different brain regions during adolescent brain development.

The observed whole-brain patterns of regional contributions are highly similar across brain fingerprint extractions for different time intervals (Fig. 3A), showing high correlations between the within-session and the two-year interval identifications (*r* = 0.949, *p* < 0.001), between the within-session and the four-year interval identifications (*r* = 0.948, *p* < 0.001), and between the two-year and four-year intervals identifications (*r* = 0.949, *p* < 0.001) (Fig. 3B). The common regions include bilateral middle temporal, superior frontal, and inferior parietal regions (Fig. 3A and Supplementary Table 1). These results indicate that the contributions of brain regions to individualized functional connectivity patterns are prominent and relatively stable over time.

**Fig. 3.**
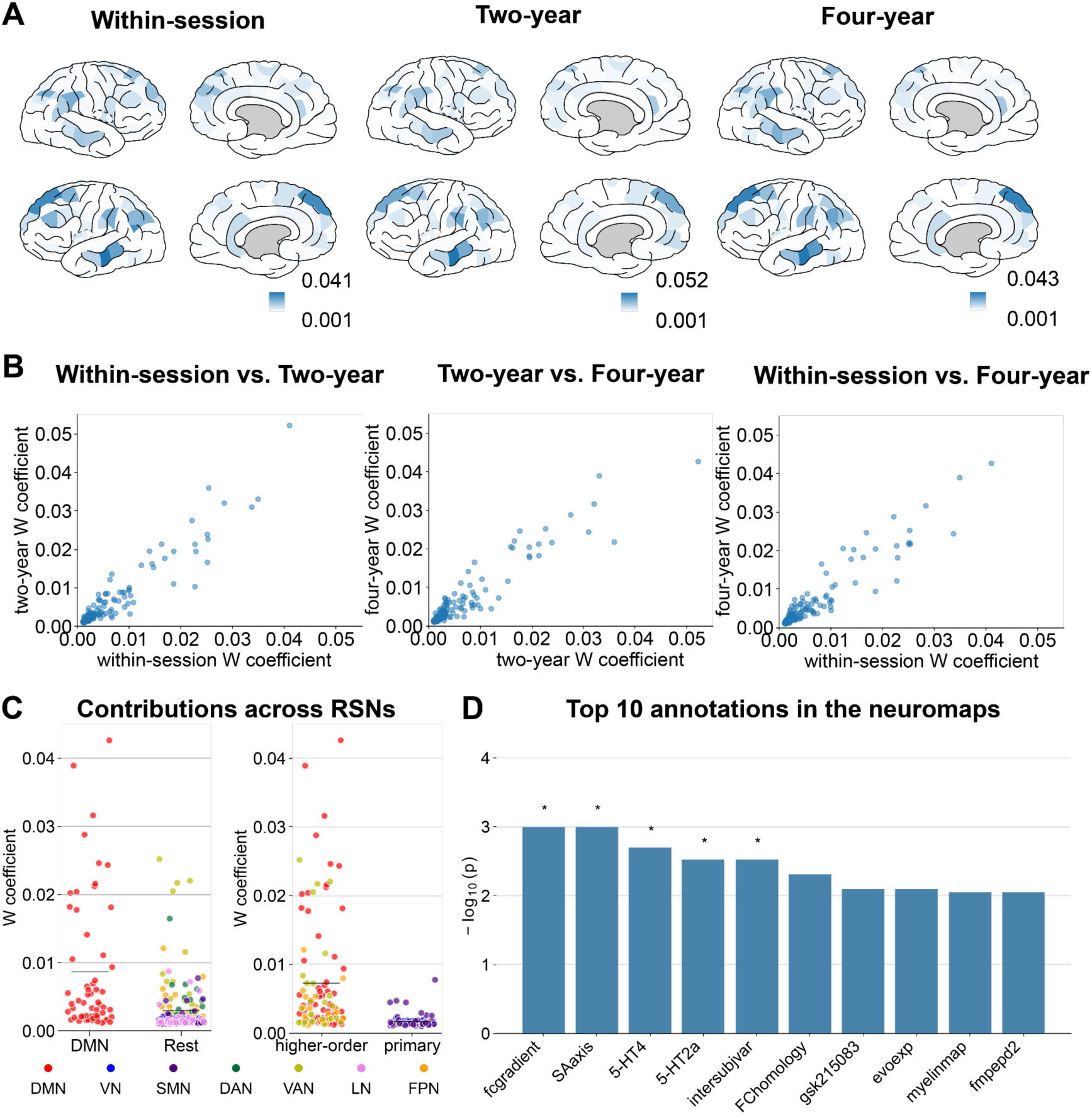
Regional contributions to the extracted brain fingerprints. (A) Brain region importance plots for time intervals of within-session, two-year, and four-year, respectively from left to right. The importance value represents the relative contribution of each region to brain fingerprint extraction. Deeper colors indicate higher importance. (B) Scatter plots demonstrate the pairwise correlation of brain region importance across three time intervals. (C) Distribution of W coefficients between the DMN and other brain networks, as well as between high-order cognitive networks and low-level motor networks. The horizontal black lines represent the mean values for each distribution. RSNs: Resting state Networks. (D) The - log(10)-transformed adjusted *p*-values for various brain networks from the Neuromaps dataset are shown. Bars marked with an asterisk (*) represent networks with statistical significance (*p* < 0.05) after FDR correction, while bars without an asterisk indicate networks with p-values above the significance threshold.

We then examined the spatial distributions of the observed regions in terms of brain functional networks (*29*). For simplicity, we report results of the brain fingerprint extraction process for four-year intervals, with results of within-session and two-year intervals shown in Supplementary Table 2&3. We found that regions from the DMN showed the highest level of the coefficient *W*, which was significantly higher than the rest of the functional networks (*t*(217) = 5.959, *p* < 0.001; Fig. 3C). The group of higher-order cognitive networks, namely the DMN, frontoparietal network (FPN), and the ventral attention network (SN), showed significantly higher levels of *W* compared to primary visual and sensorimotor networks (*t*(176) = 5.618, *p* < 0.001; Fig. 3C).

The observed patterns of regional contributions were further annotated using the neuromaps (*30*) to provide neurobiological explanations of those key regions during extracting brain fingerprints. First, considering the top ten functional gradients proposed by Margulies et al. (*31*), the observed pattern of coefficient *W* was significantly correlated to the principal gradient (*r* = 0.348, *p* = 0.001; false discovery rate (FDR) corrected *p* < 0.05 across 72 brain maps in the neuromaps; Fig. 3D), namely, the gradient from primary sensorimotor to higher-order association regions. Likewise, a similar association was observed for the sensorimotor-association (SA) axis in the brain (*r* = 0.341, *p* = 0.001; FDR corrected) (*32*). Moreover, the pattern of coefficient *W* showed a significant positive correlation (*r* = 0.320, *p* = 0.003; FDR corrected) with individual variability in functional brain connectivity in adults, as reported by Mueller 2013 (*33*), suggestive of the consistent patterns of brain functional variability between adults and early adolescents. Also, significant associations were observed for the PET tracer binding to 5-HT4 (*r* = 0.338, *p* = 0.002) and 5-HT2a (*r* = 0.269, *p* = 0.003; FDR corrected) (*34*). Additionally, the pattern of regional contributions demonstrated trend-level positive correlation with the evolutionary surface expansion from macaques to humans (*r* = 0.257, *p* = 0.008, not corrected), as well as a negative correlation with the cross-species FC homology (*r* = −0.260, *p* = 0.005, not corrected; Fig. 3D) (*35*). These results implicate the potential association between individual variability during human brain development and human-specific configurations in brain evolution (*33*, *36*).

### Cognitive correlates of the identified brain fingerprints

To investigate whether brain fingerprints generated by Metric-BolT are associated with cognitive behaviors, we used ordinary least squares (OLS) regression to examine the relationship between brain functional fingerprints and the performance on cognitive tests. Given the developmental nature of our sample, all cognitive scores were age-corrected to account for the impact of age on cognitive performance in adolescents. The OLS model revealed significant association between baseline brain fingerprints and cognitive performance for fluid intelligence (Cognition Fluid Composite, *F* = 1.282, *p* = 0.027), crystallized intelligence (Crystallized Composite, *F* = 1.405, *p* < 0.001), and executive function assessed through cardsort (Dimensional Change Card Sort, *F* = 1.290, *p* = 0.008). These results demonstrate that the identified brain fingerprints are associated with individual differences in cognitive abilities.

### The genetic influence on brain fingerprints

Having established individual brain fingerprints during development, we additionally investigated the influence of genetic factors on brain fingerprints by means of assessing the potential role of genomic relationship in brain fingerprinting. To this end, we compared the distance of brain fingerprints between subjects with a strong genomic relationship (e.g. cousins or twins, pi_hat > 0.5) and the rest of the subjects across three time intervals (see also MATERIALS AND METHODS for details). For the brain fingerprints extracted within the same session, subjects with a strong genomic relationship tended to show smaller distance of brain fingerprints as compared to the rest of the subjects (*t*(*6318*) = −12.330, *p* < 0.001; Fig. 4). Similar results were observed for the brain fingerprints extracted across the two-year interval (*t*(*1890*) = −7.467, *p* < 0.001) and four-year interval (*t*(*3780*) = −9.101, *p* < 0.001; Fig. 4). These results confirm that a strong genomic relationship is associated with more similar brain fingerprints at different developmental stages.

**Fig. 4.**
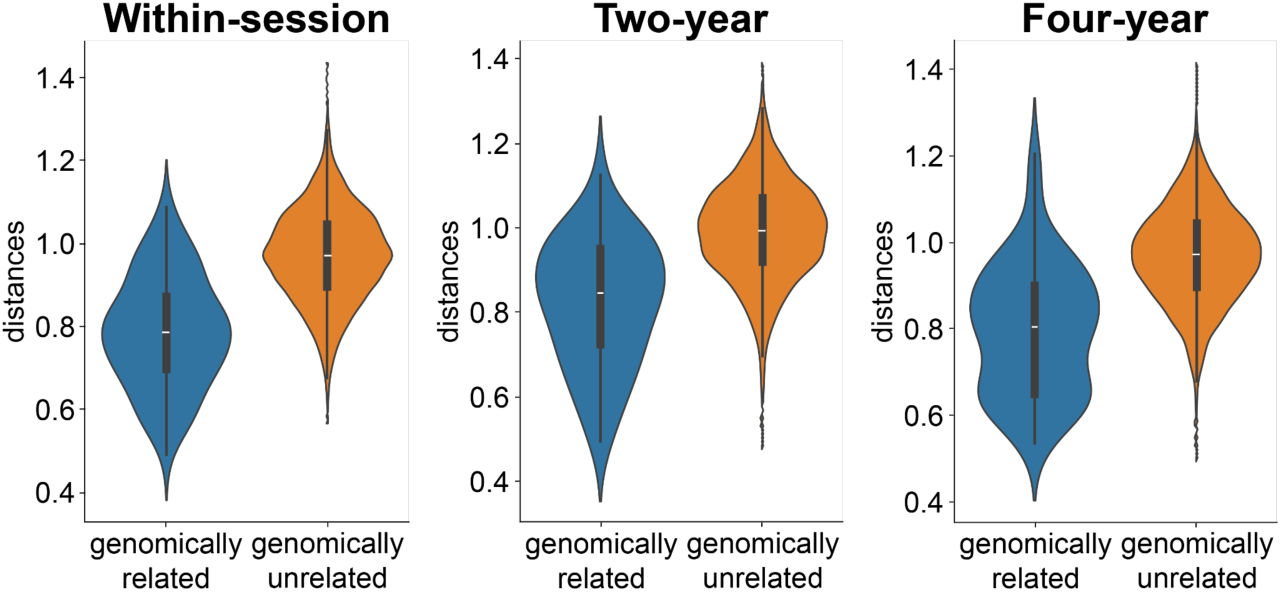
Genetic influence on brain fingerprints in development. The violin plot illustrates the distribution of distances between brain fingerprints for sample pairs, comparing those with strong genomic relationships to those without significant genomic relationships across three distinct time intervals.

## DISCUSSION

In this study we introduce a novel deep learning framework, metric-BoIT, which integrates metric learning techniques with transformer blocks to capture brain functional fingerprints during the development of early adolescents. This framework demonstrates high precision in individual identification using longitudinal fMRI data over intervals of up to four years. Brain fingerprints extracted by metric-BoIT point out the high individual variability of higher-order cognitive networks throughout development. Moreover, the brain fingerprints show significant associations with cognitive abilities, including fluid intelligence, crystallized intelligence, and executive function. These brain fingerprints are also influenced by genetic factors, suggesting a hereditary component in individual brain characteristics. Our findings demonstrate that brain functional fingerprints are effective and reliable for the developing brain, highlighting the uniqueness of brain functioning over years in early adolescents.

Our results of within-session identification point out that intrinsic brain functional fingerprints become increasingly discriminative along with age, implicating that individuals progressively become unique throughout childhood and early adolescence. This is in line with previous findings showing the emergence of individual-specific brain functional organization during neurodevelopment and the growing uniqueness of brain networks with age (*16*). This developmental trajectory prominently demonstrates that brain functional organization is a dynamic, continuous process of neural individualization, constantly adapting to unique experiences and environmental demands, and progressively forming a distinctive “brain fingerprint”. Our deep learning-based brain fingerprint identification results not only support this conclusion, but also extend this by showing a high precision of individual identification with a time span extending up to four-years, indicating a remarkable stability in the fingerprint over time. Consistent with extant literature, these findings corroborate the notion that children’s functional connectomes maintain and even enhance their self-stability throughout neurodevelopmental maturation (*37*). These findings elucidate novel explanations underlying the changes in individual stability and distinctiveness.

Brain fingerprinting during the development was noted to be driven by higher-order cognitive networks, in particular the DMN. These results are consistent with previous findings showing that higher-order association regions demonstrate substantial inter-subject variability and play crucial roles in coordinating large-scale brain activity (*38*, *39*). Additionally, the DMN not only coordinates with other brain activities but also participates in self-awareness and self-referential processing (*40*), playing an important role in development (*41*). This pattern of individual variability is known to reflect a sensorimotor-to-association axis of cortical organization (*33*, *42*), which has been suggested to arise from the temporal sequence of neurogenesis (*43–45*) and to capture the differentiated maturation rate across the cortex. Linking neurodevelopmental to evolution further revealed the pattern of individual variability to be related to evolutionary cortical expansion (*32*, *46*), which might be coordinated by the underlying gene transcription of evolutionarily important genes like the human-accelerated regions (*36*, *47*). These observations also implicate our deep-learning-based method of individual identification to be biologically explainable in terms of brain development.

Our findings further demonstrate the genetic influence on brain functional fingerprinting, demonstrating a significant difference in brain fingerprint distances between individuals with comparable and contrasting genetic backgrounds. These results align with and extend previous research, suggesting that the highly stable individual-specific characteristics may be attributed to epigenetic factors during prolonged developmental maturation (*48*, *49*). Regardless of age group, genetic inheritance significantly affects the uniqueness of functional connectivity, with stronger genomic kinship leading to more similar connectivity patterns. This indicates that genetic inheritance plays a fundamental role in shaping the uniqueness of brain functional organization from early development through to adulthood (*50*). The heritability of brain functional connectivity is primarily driven by higher-order association cortices, manifested in the FPN and DMN (*51*). Our study provides novel insights into how genetic factors shape brain functional organization during the adolescent.

Several considerations should be taken into account when interpreting our findings. First, individual identification in this study relies on resting-state fMRI data. Future work could extend brain fingerprinting to task-based fMRI data or incorporate other imaging modalities (such as EEG, Structural MRI, diffusion MRI), particularly in identifying individuals across different tasks (*52*). Secondly, unlike previous approaches using FC (*53*), the metric-BoIT model is built on time series data. While this might overlook certain information traditionally captured by FC analysis, our results suggest that this deep learning model can effectively capture complex brain dynamics (*54*, *55*) that might be missed by conventional FC-based methods. Thus, combining dynamic FC and static FC might be a promising direction to provide more comprehensive insights into studying individual differences (*56*). Finally, exploring the stability of the brain from adolescence through adulthood, is an interesting direction, as our understanding in this area remains limited. Studying the developmental trajectory of functional connectivity across these life stages can help us better understand the long-term changes and maturation of individual neural networks from youth to adulthood.

To summarize, we proposed a distance-based deep metric learning framework to provide an intuitive and interpretable approach for brain fingerprinting and dissecting inter-individual variability in functional dynamics in early adolescents. This method achieved a high precision in individual identification over as long as four years and pointed to the contribution of higher-order association networks in fingerprinting. The extracted brain fingerprint exhibits the association with behavior and is modulated by genetic factors. Our proposed approach provides new insights into the unique brain developmental trajectory in early adolescents.

## MATERIALS AND METHODS

### MRI Data

MRI data were obtained from the ABCD Study (data release 5.1; DAR ID: 16920), which includes longitudinal MRI from 11,892 nine to ten-year-old children (*57–61*). These participants and their families were recruited through schools and community settings across 21 sites in the United States, representing a diverse cross-section of the U.S. population in terms of education, race, and socioeconomic status. The ethical review of the study was approved by the University of California’s central Institutional Review Board (IRB) for most sites, while local IRBs approved the study at other locations. The current study used data of 1,325 participants with an initial age range of 9 to 10 years, excluding all subjects without the fourth-year follow-up data and those without either T1 imaging or resting-state fMRI data (Supplementary Fig. 1).

### Data preprocessing

Minimally processed T1 weighted MRI data and resting-state fMRI data were obtained from the ABCD study. Details of scanning parameters and processing procedures have been reported in (*62*). Minimally-processed T1-weighted MRI data were further processed using FreeSurfer (v.6.0) for brain segmentation and cortical mantle reconstruction. The FreeSurfer output was manually inspected by experienced researchers. The reconstructed cortical mantle was parcellated into 219 distinct regions according to a subdivision of the Desikan–Killiany (DK-219) atlas (*63*, *64*).

In parallel, minimally-processed resting-state fMRI data were further processed using CATO (v.3.1.2) following the following steps (*65*). First, resting-state fMRI data were first realigned using MCFLIRT to correct for any displacement during participant scanning. Second, data were co-registered with the T1-weighted data to overlap with the resultant cortical parcellation maps under the DK-219 atlas. Then, six motion parameters, first order drifts of the six motion parameters, and mean signal intensity of voxels in white matter and CSF were regressed out through linear models. Bandpass filtering was further conducted to eliminate frequencies outside the 0.01 to 0.1 Hz range, retaining only signals within the desired frequency band. Notably, scrubbing was not applied to ensure the same length of time series across participants.

### Metric-BolT model construction

We proposed a distance-based metric learning method to make the brain fingerprints extracted from different time series from the same subject more proximate, and those from different subjects more distinct (Fig. 1). Specifically, each participant is treated as a class, and the matrix of fMRI time series of each participant is treated as a sample. The deep learning model then maps each sample (i.e. time series) into a high-dimensional space. In this space, the ideal goal is to achieve a configuration in which the maximum intraclass distance is always smaller than the minimum interclass distance. This means the data from different subjects are well separated, while the data from the same subject are closely clustered together. This approach consists of two key steps: (1) encoding fMRI time series into feature vectors, referred to as brain fingerprints, and (2) comparing the distance of feature vector pairs. In the first step, we employed the BolT model (*28*) to optimally represent the fMRI data in the embedding space, resulting in a highly representative encoding of the original fMRI time series (see for details in Supplementary Methods). In the second step, We quantified the distance between feature vectors using 1 minus cosine similarity and applied the cosine similarity measure in TripletMarginLoss as loss function (*66*), which is defined as:

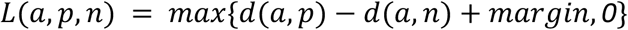

*L* represents the triplet loss function, where *a*, *p*, and *n* denote the anchor, positive sample (same class as *a*), and negative sample (different class from *a*), respectively. The *margin* is a threshold used to control the distance between positive and negative samples. This loss function pushes samples of the same class closer and samples of different classes farther apart, as illustrated in Fig. 1. This two-step process allows us to establish brain fingerprints for each individual and effectively distinguish between different individuals in the learned embedding space. The whole dataset was randomly divided into training (80%), validation (10%), and test (10%) sets. This random splitting was applied independently to each time point and time interval.

### Individual identification

To validate the effectiveness of the trained model, specifically the uniqueness and stability of brain fingerprints, we conducted individual identification experiments across six cases: two runs from three within-session and three cross-session longitudinal cases on the test set. Prior to the identification process, all time series from both runs were processed through Metric BolT to generate brain fingerprints for each subject. Then the feature vectors from one run were used to create the brain fingerprint database, *D* = [*X_i_*, *i* = *1*, . . ., *N*], where *i* stands for the participant, *N* is the number of subjects in the test set. Feature vectors from the other run were used as the target for identification. In each iteration, a target brain fingerprint was matched to the database of fingerprints for *N* iterations, the identity of a target feature vector *Y_i_* was predicted by comparing it to all vectors in *D*. The predicted identity was determined by finding the vector in *D* with the minimal distance to *Y_i_*. Note that we performed identification with replacement, meaning each target feature vector could be matched to any subject in the database, regardless of previous matches. This process was repeated for all feature vectors in the target run. To ensure robust evaluation, we employed a bi-directional testing strategy: we swapped the target and database run and repeated the entire procedure.

Three key metrics were established to evaluate the efficacy of our experimental approach: success rate (SR), intra-class to inter-class distance ratio (CR), and silhouette coefficient (SC) (see for details in Supplementary Methods). To assess the statistical significance of the identification accuracy, we used a nonparametric permutation test. This approach involved 1,000 independent tests, where in each test, subject identities in the target set were randomly permuted. After each randomization, the identification algorithm was applied, and the SR was recorded. This process generated a null distribution of SR scores. By comparing the originally observed SR to this null distribution, we determined the statistical significance of the results, evaluating the method’s performance beyond chance level.

### Brain fingerprint interpretation

We calculated the contribution of each brain region to gain a deeper understanding of the underlying brain functional dynamics that drive brain fingerprinting. We first used an attention mechanism-based approach to quantify the contribution of each BOLD token to the brain fingerprint, resulting in token-level importance for each participant (Supplementary Methods). Based on this, we used a random forest approach to obtain the importance of brain regions in fingerprinting, translating these token-level importance into brain region importance. By labeling the top five most important BOLD tokens as 1 and the top five least important as 0, we created a binary classification task. The resulting model weights, *W*, offered a quantitative measure of each brain region contribution to brain fingerprint.

The cortical patterns of regional contributions were further annotated in terms of the seven resting-state brain functional networks (*29*) and distinct biological brain maps. Here, we utilized the Neuromap toolkit to conduct a correlation analysis between these brain region contributions and multidimensional brain maps (*30*). The specific process involved mapping 219 brain regions to the fsaverage standard space, downsampling both the built-in Neuromaps and the standard space brain maps to fsaverage ‘3k’, then calculating Pearson correlations, and performing spatial statistical correction through 1000 spin tests (*67*). Using the correspondence between nodes in the Lausanne250 brain atlas and seven resting-state functional networks provided by neuromaps, we performed *t*-tests comparing each network to all other networks, as well as *t*-tests comparing higher-order cognitive networks to lower-order sensorimotor networks.

### Correlation analysis between brain fingerprints and cognitive abilities

To examine the association between brain fingerprints and cognitive abilities, we employed least squares regression and tested the significance of regression coefficients. Specifically, we selected measures of higher-order cognition, including fluid intelligence (Cognition Fluid Composite, mean ± SD: 96.944 ± 17.685, range: 58– 138), crystallized intelligence (Crystallized Composite, mean ± SD: 108.252 ± 20.932, range: 36–165), executive function (Dimensional Change Card Sort, mean ± SD: 98.330 ± 16.578, range: 77–155), and general cognitive ability (mean ± SD: 102.832 ± 19.269, range: 59–154) from the (*68*). Given the substantial individual differences in neural development during childhood and adolescence, and the non-linear effects of age on cognitive performance, we utilized age-corrected cognitive scores to eliminate potential assessment bias caused by varying developmental rates, thereby more accurately reflecting individual cognitive capabilities. We then constructed a linear regression model with brain fingerprint vectors as independent variables and cognitive test scores as the dependent variable. After fitting the model using the least squares method, we proposed a null hypothesis that all regression coefficients equal zero (indicating no association between brain fingerprints and cognitive scores). We then calculated the *F*-statistic and its corresponding *p*-value for the regression coefficients to assess whether brain fingerprints could significantly explain the variance in cognitive abilities.

### Genetic influence on brain fingerprints

We examined whether genetic factors affect the extracted brain fingerprints. We used genotype data and computed the Identity by Descent (IBD) measure, represented by the pi_hat statistic. IBD refers to the inheritance of identical genetic material from a common ancestor (Supplementary Methods) (*69*), with the genomic relationship quantified by pi_hat. Pi_hat ranges from 0 to 1, with values above 0.5 typically indicating first-degree kinship. We conducted experiments across three time-intervals through the following steps: we categorized the subjects into two groups based on their pi_hat values, with pi_hat > 0.5 defining the group exhibiting strong genomic relationship and pi_hat ≤ 0.5 representing genetically unrelated. We created new training, validation, and test sets using only genomically unrelated participants. Training then proceeded on this newly defined training set to develop a new metric-BolT. Subsequently, we tested the trained model on pairs of genetically similar subjects and pairs of genetically dissimilar subjects, comprehensively assessing the genetic specificity of brain fingerprints. The number of subjects with close genetic relationships and the rest of the subjects for the within-session, two-year, and four-year intervals were 80 and 6240, 44 and 1848, and 62 and 3720, respectively. Finally, we used two sample t-tests to compare group means of distances. A *p*-value of < 0.05 was considered statistically significant across all tests.

## ACKNOWLEDGEMENTS

This study received funding from the National Natural Science Foundation of China (grant number 82202264 to Y.W. and grant number 82301734 to T.Q.), Beijing Municipal Natural Science Foundation (grant number 7232341 to Y.W.), Analyses were supported by the High-Performance Computing Platform of BUPT.

Data used in the preparation of this article were obtained from the Adolescent Brain Cognitive Development (ABCD) Study (https://abcdstudy.org), held in the NIMH Data Archive (NDA). This is a multisite, longitudinal study designed to recruit more than 10,000 children aged 9-10 and follow them over 10 years into early adulthood. The ABCD Study® is supported by the National Institutes of Health and additional federal partners under award numbers U01DA041048, U01DA050989, U01DA051016, U01DA041022, U01DA051018, U01DA051037, U01DA050987, U01DA041174, U01DA041106, U01DA041117, U01DA041028, U01DA041134, U01DA050988, U01DA051039, U01DA041156, U01DA041025, U01DA041120, U01DA051038, U01DA041148, U01DA041093, U01DA041089, U24DA041123, U24DA041147. A full list of supporters is available at https://abcdstudy.org/federal-partners.html. A listing of participating sites and a complete listing of the study investigators can be found at https://abcdstudy.org/consortium_members/. ABCD consortium investigators designed and implemented the study and/or provided data but did not necessarily participate in the analysis or writing of this report. This manuscript reflects the views of the authors and may not reflect the opinions or views of the NIH or ABCD consortium investigators.

## AUTHOR CONTRIBUTIONS

R.X. developed the model and performed the analyses. Y.W. and T.Q. conceived the idea of this study. Y.W. and Y.L. supervised this study. S.Z. performed functional annotation analysis. Q.J., Z.W., K.Z., S.Z, J.Z., Y.L. provided critical revision with important intellectual content. R.X., T.Q., and Y.W. wrote the manuscript with contributions from all coauthors.

