## Supplementary Materials for "Functional fingerprinting for the developing brain using deep metric learning: an ABCD study"

### Supplementary Methods

#### Metric-BolT implementation details

For encoding time series into feature vectors (i.e.brain fingerprint), we employed the Blood-oxygen-level-dependent Transformer (BolT) model (*1*) to optimally represent the fMRI data from the ABCD dataset in the embedding space. Based on the transformer architecture (*2*), each time point's BOLD response in the time series is treated as a BOLD token, and BolT can directly operate on these BOLD tokens. To capture local representations, BolT splits the time series into temporally-overlapping windows and employs a cascade of transformer blocks to encode window-specific representations of BOLD tokens. To enhance expressiveness across broad time scales without elevating computational costs, BolT leverages a novel fused window attention mechanism that utilizes cross attention and token fusion among overlapping windows. While cross attention enables interactions between base BOLD tokens in a given window and fringe tokens in neighboring windows prior to encoding, token fusion enables integration of encoded representations across neighboring windows. To hierarchically transition from local to global representations, the extent of window overlap in transformer blocks is progressively increased across the cascade. BolT improves performance by utilizing classification (CLS) tokens to capture high-level features (*3*). Window-specific CLS tokens are introduced to maintain local sensitivity and compatibility with the hierarchical model structure. Meanwhile, information exchange is promoted by a novel cross-window regularization that aligns these CLS tokens across windows. At the end of the cascade, the encoded CLS tokens are averaged across windows. This averaging process consolidates both local and global information, resulting in a highly representative encoding of the original time series.

For comparing feature vector pairs for distance, we focused on two critical components: the distance measure and the loss function. Given that our Metric-BolT model maps the original input into a high-dimensional vector space, we use cosine similarity to measure the similarity between two vectors and use 1 minus cosine similarity as the distance measure. Cosine similarity is computationally efficient in high-dimensional spaces, with values ranging from -1 to 1, where higher values indicate greater similarity between vectors. For the loss function, we employed TripletMarginLoss (*4*), which outperformed alternatives such as MultiSimilarityLoss (*5*), CircleLoss (*6*) in terms of efficiency and effectiveness in our experiment. The TripletMarginLoss is defined as:

$$L(a,p,n) = max\{d(a,p) - d(a,n) + margin, 0\}$$

$L$ represents the triplet loss function, where $a$, $p$, and $n$ denote the anchor, positive sample (same class as $a$), and negative sample (different class from $a$), respectively. When $d(a,p) - d(a,n) + margin > 0$, indicating the negative sample is not sufficiently far from the anchor compared to the positive sample, this difference is recorded as the loss. Otherwise, the loss is set to 0. This mechanism encourages the model to learn that the distance between anchor-positive pairs is smaller than anchor-negative pairs by at least the margin, thus pushing similar samples closer and dissimilar samples farther apart, as illustrated in Fig. 1.

**Evaluation metrics**

The success rate was defined as the proportion of subjects whose identity was accurately identified:

$$SR = \frac{number\ of\ correctly\ identified\ subjects}{number\ of\ total\ subjects}$$

Correct identification was scored as 1 point for the correct match (identified identity matches the true identity) and 0 points for an incorrect match. Given that our metric learning shares conceptual similarities with clustering, we employed two additional clustering metrics to provide a more comprehensive analysis of the individual identification performance. For this, each subject was treated as a class, with their corresponding feature vectors serving as samples. The intra-class to inter-class distance ratio was utilized to assess the compactness within classes relative to the separation between classes. The ratio is calculated as:

$$CR = \frac{average\ intra - class\ distance}{average\ inter - class\ distance}$$

where the intra-class distance is the average distance between samples within the same class, while the inter-class distance is the average distance between samples from different classes. The CR ranges from 0 to 1, with smaller values indicating that samples within a class are more compact and better separated from those in other classes. Complementing this, the silhouette coefficient was employed to quantify the cohesion and separation of the resulting clusters. The silhouette coefficient for a sample i is defined as:

$$SC(i) = \frac{b(i) - a(i)}{max(a(i), b(i))}$$

$$SC = \frac{1}{N}\sum_{i=1}^{N} SC(i)$$

where $a(i)$ is the mean distance between $i$ and all other vectors in the same subject, and $b(i)$ is the mean distance between $i$ and all points in the nearest neighboring subject. The overall silhouette coefficient is the average across all subjects and ranges from -1 to 1, where: values near 1 indicate that the vector is well-matched to its own subject and poorly-matched to neighboring subjects. Values around 0 suggest the vector lies near the boundary between two subjects. Negative values indicate that the vector might have been misclassified. This comprehensive evaluation framework allows for a rigorous assessment of the method's performance from multiple perspectives, enhancing the robustness of our findings.

**Model interpretations using the transformer block**

For each transformer block, we first compute the attention map for each window:

$$\bar{A}_{mi} = E_h((A_i)^+)$$

where $m$ denotes the index of the transformer block, $i$ denotes the index of the window, $E_h$ denotes the averaging operator across attention heads for aggregation,+ denotes rectification to prevent negative values, and then aggregate the attention matrix for all time windows for each transformer block to form a global attention graph $\bar{A}_G$, ($\bar{A}_G \in R^{(F+T)*(F+T)}$, where $F$ denotes the number of time windows and $T$ denotes the number of BOLD tokens). The values in $\bar{A}_G$ represent the attention between each token. Specially, the $\bar{A}_G[:F, F:]$ represents the attention weights from the CLS tokens to the BOLD tokens. This can be interpreted as the contribution or influence of the BOLD tokens on the CLS tokens.

Next, a token-relevance matrix $Rel[0]$ is initialized to represent the interactions between each token in the whole procedure of extracting brain fingerprints, which is initialized as an identity matrix of size $(F+T)*(F+T)$, indicating that each token has only autocorrelation, and then the token correlation maps of each transformer block are updated step by step using the attention matrix :

$$Rel[m+1] = Rel[m] + \bar{A}_G[m]Rel[m]$$

Following the calculation of the token-relevancy map, importance weights for input BOLD tokens are finally derived as:

$$w_{imp} = \frac{1}{F}\sum_{i=0}^{F-1} Rel[M](i, F:)$$

The final importance weight of each BOLD token was derived from its cross-window average correlation with the CLS token, providing a measure of its overall contribution to the brain fingerprint.

**Genetic data analysis**

In the process of genotype data quality control, we first calculate the missing rates for samples and Single Nucleotide Polymorphisms (SNPs) to exclude low-quality data. Next, we calculate the minor allele frequency (MAF), which helps in categorizing and analyzing the frequency of variants. Then, through the Hardy-Weinberg equilibrium exact test, we can detect potential genotyping errors. To ensure data accuracy, we apply multiple filters to exclude samples and SNPs that do not meet standards. Meanwhile, through linkage disequilibrium (LD) pruning, we effectively remove SNPs with strong linkage disequilibrium, reducing redundancy in the analysis. Additionally, we check the heterozygosity coefficient of samples, using the inbreeding coefficient to identify and exclude potentially contaminated samples. Finally, we select specific samples and SNPs for further analysis to enhance the reliability and accuracy of the data.

By calculating IBD (the proportion of shared alleles) and evaluating PI_HAT (the kinship coefficient), we assess the genetic relationships between individuals. IBD refers to the situation where two individuals share identical alleles and these alleles are inherited from a common ancestor rather than arising from mutation or other means. PI_HAT is a statistical measure used to quantify the concept of IBD, calculated using the followed formula:

$$PI\_HAT = P(IBD = 2) + 0.5 * P(IBD = 1)$$

where $P(IBD = 2)$ represents the probability that both alleles come from a common ancestor, while $P(IBD = 1)$ signifies the probability that only one allele comes from a common ancestor. The value of $PI\_HAT$ ranges from 0 to 1, reflecting the extent to which two individuals share alleles.

**Supplementary Figures**

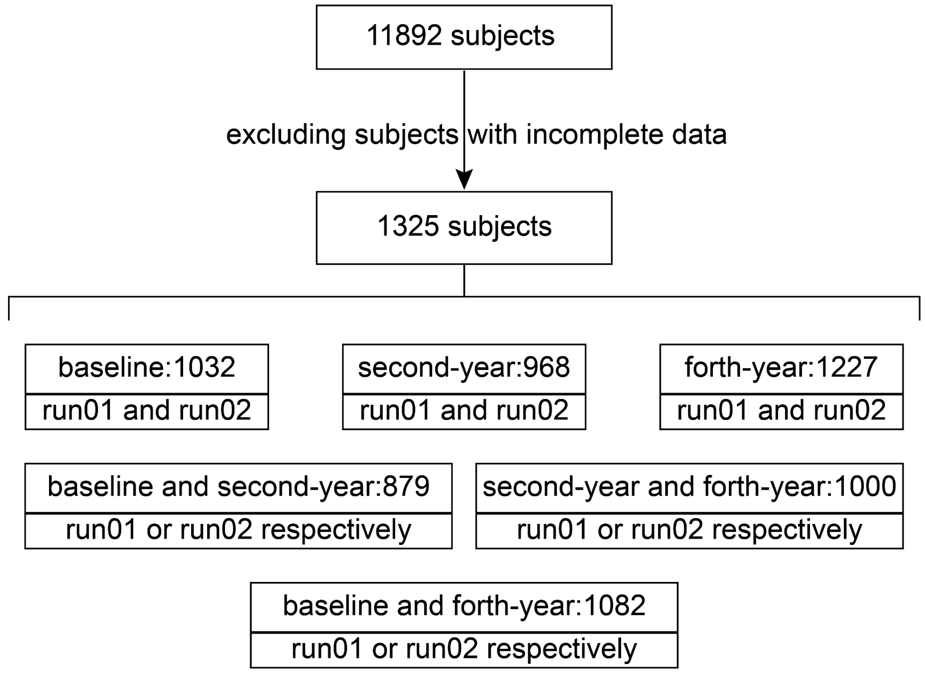


**Supplementary Fig. 1 Participant Screening Pipeline.** Subjects with incomplete data were defined as those without the fourth-year follow-up data or those lacking either T1 imaging or resting-state fMRI data. "run01 and run02" denotes the availability of both runs at the specified time point. "run01 or run02 respectively" signifies that at each of two distinct time points, at least one of the two runs (run01 or run02) is available.

**Supplementary Tables**

**Table S1. The top five most important brain regions and their W coefficient across three time spans.**

| within-session | | two-year | | four-year | |
|---|---|---|---|---|---|
| region | *W* coff | region | *W* coff | region | *W* coff |
| lh-middletemporal_3 | 0.041 | lh-middletemporal_3 | 0.052 | lh-middletemporal_3 | 0.043 |
| lh-superiorfrontal_3 | 0.035 | lh-supramarginal_3 | 0.036 | lh-superiorfrontal_3 | 0.039 |
| lh-inferiorparietal_3 | 0.034 | lh-superiorfrontal_3 | 0.033 | lh-middletemporal_2 | 0.032 |
| lh-middletemporal_2 | 0.028 | lh-middletemporal_2 | 0.032 | lh-inferiorparietal_5 | 0.029 |
| lh-supramarginal_3 | 0.025 | lh-inferiorparietal_3 | 0.031 | rh-supramarginal_3 | 0.025 |

**Table S2. Contribution of brain networks extracting brain fingerprints over a within-session and two-year span.**

| Brain map | within-session | | two-year | |
|---|---|---|---|---|
| | r-value | p-value | r-value | p-value |
| fcgradient | 0.387 | 0.001 | 0.348 | 0.001 |
| SAaxis | 0.370 | 0.001 | 0.341 | 0.001 |
| 5-HT4 | 0.323 | 0.008 | 0.338 | 0.002 |
| 5-HT2a | 0.261 | 0.007 | 0.269 | 0.003 |
| intersubjvar | 0.350 | 0.004 | 0.320 | 0.003 |
| FChomology | -0.288 | 0.004 | -0.260 | 0.005 |
| gsk215083 | 0.189 | 0.023 | 0.204 | 0.008 |
| evoexp | 0.260 | 0.009 | 0.257 | 0.008 |
| myelinmap | -0.254 | 0.016 | -0.242 | 0.009 |
| fmpepd2 | 0.287 | 0.005 | 0.260 | 0.009 |

**Table 3. T-values and p-values from the t-test of W-values and brain networks over a two-year and within-session span.**

| Networks | within-session | | two-year | |
|---|---|---|---|---|
| | t-value | p-value | t-value | p-value |
| Visual Network | -3.076 | 0.002 | -2.726 | 0.007 |
| Sensorimotor Network | -3.042 | 0.003 | -2.647 | 0.009 |
| Dorsal Attention Network | -0.557 | 0.578 | -0.320 | 0.749 |
| Ventral Attention Network | 1.688 | 0.093 | 1.523 | 0.129 |
| Limbic Network | -2.226 | 0.027 | -1.751 | 0.081 |
| Fronto-parietal Network | 0.334 | 0.739 | -0.624 | 0.533 |
| Default Mode Network | 6.021 | <0.001 | 5.517 | <0.001 |

**Table 4. Statistical results of associations between brain fingerprints and cognitive behaviors obtained from NIH Toolbox.**

| Cognitive abilities | F-value | p-value |
|---|---|---|
| Cognition Fluid Composite | 1.228 | 0.027 |
| Crystallized Composite | 1.521 | <0.001 |
| Dimensional Change Card Sort | 1.290 | 0.008 |
| Cognition Total Composite | 1.503 | <0.001 |
| Oral Reading Recognition | 1.242 | 0.020 |
| Picture Vocabulary | 1.469 | <0.001 |
| List Sorting Working Memory Test | 1.194 | 0.047 |
| Flanker Inhibitory Control and Attention Test | 1.109 | 0.164 |
| Pattern Comparison Processing Speed Test | 1.101 | 0.181 |
| Picture Sequence Memory Test | 0.930 | 0.741 |
